# Female sex hormones enhance gonococcal colonization at the endocervix by modifying cervical mucus

**DOI:** 10.1101/2025.08.14.670271

**Authors:** Sofia Di Benigno, Vonetta Edwards, Shrestha Mathur, Allison Boboltz, Elizabeth M. Engle, Logan Kaler, Margaret A. Scull, Gregg Duncan, Liang-Chun Wang, Daniel C. Stein, Wenxia Song

**Affiliations:** Department of Cell Biology & Molecular Genetics, University of Maryland, College Park, MD 20742, USA; Institute for Genome Sciences University of Maryland School of Medicine, Baltimore, MD 21201, USA; Department of Microbiology and Immunology, University of Maryland School of Medicine, Baltimore, MD, 21201, USA; Fischell Department of Bioengineering, University of Maryland, College Park, MD 20742, USA; Marine & Pathogenic Microbiology Lab, National Sun Yat-Sen University, Kaohsiung, Taiwan

**Keywords:** Neisseria gonorrhoeae, Colonization, Sex hormones, Mucus, Human cervix

## Abstract

*Neisseria gonorrhoeae* is a human-exclusive pathogen that causes gonorrhea. Gonococci (GC) initiate female infections by colonizing the cervix, which can remain asymptomatic, cause cervicitis, or ascend to the upper female reproductive tract (FRT), leading to severe tissue damage. The FRT undergoes sex hormone-mediated changes during the menstrual cycle, which have long been implicated in the vulnerability to GC infection. One of the major changes is the increase and decrease in the production of the gel-forming mucin MUC5B by the endocervix in response to the level of estradiol (E2). This study examined the impact of sex hormones on GC infection of the human cervix, utilizing a human cervical tissue explant model. Tissue explants were treated without and with E2 alone or in combination with progesterone (E2+P4) to mimic various menstrual cycle phases. Treatment of E2 or E2+P4 enhanced GC colonization at the endocervix exclusively, but did not affect epithelial transmigration. While both treatments increased the number of GC microcolonies, E2+P4 increased GC colony size on the endocervical epithelium. These increases were independent of GC host receptors, carcinoembryonic antigen-related cell adhesion molecules. GC effectively diffused through cervical mucus to interact with the cervical epithelium under all hormone conditions and through mucin hydrogels with different MUC5B and MUC5AC compositions. Mucus gels collected from cervical explants and animal mucin mixtures enhanced GC aggregation *in vitro*. GC diffusion through mucin-hydrogels and aggregation in the presence of cervical mucus or animal mucins decreased as the MUC5B concentration increased. Our results suggest that female sex hormones promote GC colonization at the human endocervix by modulating the cervical mucus production, regulating women’s susceptibility to GC infection, and further reveal the ability of GC to evade the mucus defense barrier for infection.

**Author Summary:** *Neisseria gonorrhoeae* is a bacterial pathogen that primarily infects the human genital and female reproductive tracts, causing gonorrhea. While this bacterium can infect both men and women, the infection can lead to severe and permanent damage to women’s reproductive systems. Currently, the relationship of gonococcal infection with the menstrual cycle is unknown. Here, we utilize a human cervical tissue explant model that mimics gonococcal infection in women to examine the impact of female sex hormones that drive the menstrual cycle on gonococcal infection. We found that estrogen alone or in combination with progesterone enhanced gonococcal colonization, increasing both the number and size of bacterial microcolonies on the cervical luminal surface, through regulating mucus production. Gonococci effectively penetrate through mucus layers to reach cervical epithelial cells and also prefer to aggregate with each other in the presence of mucus. Our results reveal that hormone-regulated mucus production changes the vulnerability of women to gonococcal infection, and that gonococci convert the mucus defense barrier into a colonization facilitator.

## Introduction

*Neisseria gonorrhoeae* (GC) is a human-exclusive pathogen and is the etiologic agent of the second most common sexually transmitted bacterial infection. It poses a serious threat to public health due to increasing antibiotic resistance and frequent asymptomatic infections. It has been estimated that more than 50% of infections in both male and female individuals do not present with noticeable symptoms [1, 2]. Consequently, infections remain untreated and result in silent transmission and serious sequelae [2, 3]. Female patients bear the brunt of these sequelae, with ∼10-20% of untreated infections ascending from the lower to the upper female reproductive tract (FRT), leading to pelvic inflammatory disease (PID) [1, 4, 5]. The resulting inflammation can scar the uterus and/or fallopian tubes, causing lifelong pain and infertility and increasing the risk for life-threatening ectopic pregnancy [4, 6]. Despite the unique risks to female patients, there remains little understanding of how GC causes such a wide variety of clinical presentations in the FRT.

The cervix is the primary site of GC colonization [7, 8] for both symptomatic and asymptomatic infections and also the only path for GC to ascend from the vagina to the uterus. The cervical epithelia transition from flat, multi-layered squamous epithelial cells on the ectocervix to a single layer of highly polarized columnar epithelial cells on the endocervix [9]. Colonization is initiated by pili attachment to the luminal surface of the epithelium [10]. Previous studies using either primary and immortalized human cervical epithelial cells [11, 12] or a human cervical tissue explant model [13] have shown that pili are essential for GC infection. Pili retraction allows GC to interact more intimately with the epithelium through GC surface molecules [14, 15], such as opacity-associated proteins (Opa) and lipooligosaccharides (LOS), securing colonization [16–18].

GC colonization can induce the disassembly of cervical epithelial cell-cell junctions, which leads to the shedding of GC-colonized epithelial cells, reducing colonization [13, 19]. GC-induced apical junction disassembly of the monolayered columnar endocervical epithelium provides opportunities for GC to penetrate the epithelium into the subepithelium, promoting invasive and disseminated infection [13, 20]. Opa proteins, a family of major surface proteins, primarily interact with human carcinoembryonic antigen-related cell adhesion molecules (CEACAMs) on the epithelial surface [21]. Opa-CEACAM interactions inhibit epithelial cell-cell junction disassembly, which prevents GC-associated epithelial cells from shedding, promotes colonization, and also reduces GC penetration of the endocervical epithelium into the subepithelium [13, 20].

A distinct feature of the FRT, the site of GC infection in females, is the menstrual cycle, driven by the cyclic changes of the sex hormones estradiol (E2) and progesterone (P4) [22, 23]. The follicular phase is characterized by a steady rise in E2 levels during the first 14 days of the cycle, which facilitates the development and maturation of an ovarian follicle. E2 levels reach a sustained peak for at least 48 hours at the end of the follicular phase, which triggers a surge in luteinizing hormone and follicle-stimulating hormone. This gonadotropin surge induces ovulation and a rapid decline in E2 levels. During the luteal phase, the ovarian corpus luteum develops and secretes high levels of P4, peaking for two days. P4, along with a smaller secondary peak of E2, dominates the luteal phase. When P4 levels decrease, menses are initiated. The levels of these four hormones in the blood and their functions in the reproductive tract are well-established, with the E2 peak levels at ∼350-500 pg/mL at the follicular phase before ovulation and ∼200 pg/mL at the luteal phase and the P4 peak level at 10 ng/mL at the Luteal phase [24]. The levels of hormones in cervical tissue have not been well-studied; however, they are likely much higher than blood levels due to the proximity of the cervix to the ovaries, where local E2 and P4 secretion occurs. Further, whether and how the hormonal cycle influences GC infection remains elusive.

Clinical studies have found that the days surrounding ovulation have been suggested to be “a window of vulnerability” in the FRT [25, 26]. This window is associated with a decreased immune response, which is likely necessary for tolerating sperm while providing potential pathogens an opportunity for infection. HIV infections have been reported to be more likely in the week immediately following ovulation [25, 27, 28]. However, GC were more likely to be recovered from a cervical swab of a patient presenting at a clinic during the first five days of menses [29], and GC recovered during menses were also more likely to be Opa-free [30]. Furthermore, individuals who were exposed to GC through intercourse during menses were also more likely to produce a positive cervical swab when they later presented at a clinic, regardless of what stage of the cycle they were in at the time of presentation [31]. On the other hand, oral contraceptives, which are usually composed of ethinyl estradiol and progestin, were suggested to be protective, probably by preventing the onset of menses [32]. These data support that the hormonal changes during the menstrual cycle regulate the interactions of the FRT with various pathogens. However, the mechanisms underlying such impacts remain unknown.

The impact of the sex hormonal cycle on epithelial shedding and tissue reorganization of the endometrium and the uterus is well known [9, 33, 34], but its effects on the human cervix remain unclear. *In vitro* studies have shown that E2 treatment of primary cervical epithelial cells increased the paracellular permeability of the tight junction [9, 35, 36]. This potentially allows for increased fluid transport to promote movement of mucus and spermatozoa through the FRT [37, 38]. It is still unknown, however, whether and how these hormonal effects on cervical epithelial cell-cell junctions impact GC infection.

The hormonal cycle induces a cyclic change in the mucus production of the human cervix to facilitate sperm swimming to the upper FRT during ovulation and safeguard pregnancy from infection [39, 40]. The mucus consists of two forms, a membrane-anchored mucin layer and an overlying gel-forming layer [41]. The physical and chemical properties of the cervical mucus can be regulated by the type of gel-forming mucins expressed [42]. The primary gel-forming mucins in the cervical mucus are MUC5B and MUC5AC, with MUC6 expressed at much lower levels and MUC2 expressed intermittently [39, 43]. These mucins form gels through a combination of disulfide bonds, hydrophobic interactions, and electrostatic repulsion to form a protective mesh over the luminal surface of the epithelium [44, 45]. MUC5B-dominated mucus at sites such as the airway is a viscoelastic gel and primarily facilitates mucociliary transport while providing a physical barrier [46, 47], while MUC5AC-dominated mucus in the GI tract creates a stiffer gel and plays a protective function [48, 49]. In the cervix, E2 and P4 levels dictate the amount and ratio of MUC5B and MUC5AC expressed, consequently impacting its physical properties. MUC5B production in the endocervix increases at midcycle when the E2 level is high, dominating the gel-forming layer [50, 51], which reduces the viscoelasticity of the mucus and promotes sperm movement from the lower to the upper FRT [40]. MUC5B expression decreases as the P4 level rises, which increases mucus gel stiffness, providing a physical barrier [40, 43, 50, 51]. Mucus barrier changes during health and disease in airway have been studied [52, 53], whether and how the cyclic changes of the cervical mucus influence GC infection are unknown.

This study examined how the female sex hormones E2 and P4 modulate GC infectivity in the human cervix, using a human cervical tissue explant model that we previously established and a MS11 WT GC strain expressing phase-variable pili and Opa proteins. Our results show that a 72-h treatment with E2 alone or in combination with P4 increases GC colonization on the human endocervix exclusively. E2 alone increases the number of GC microcolonies on the epithelial surface, while the combination of E2 and P4 has the additional effect of increasing the size of GC microcolonies. However, the hormone treatment did not increase the expression of GC host receptor CEACAMs nor did it affect epithelial cell-cell junctions and epithelial shedding. Interestingly, GC efficiently penetrated the mucus layer on the cervical epithelium of tissue explants and animal mucin hydrogels with various MUC5B and MUC5AC ratios. Furthermore, both the cervical mucus collected from the tissue explants and commercially available animal mucin proteins significantly increased GC aggregation *in vitro*. Our data suggest that sex hormones enhance GC colonization at the human cervix by modifying the composition of secreted mucus, revealing a new mechanism for GC adapting to the ever-changing environment of the FRT for survival.

## Results

### Estradiol and progesterone treatments enhance GC colonization at the endocervix by increasing GC colony number and size

To examine the impact of sex hormones on GC infectivity in the human cervix, we utilized a human cervical tissue explant model that has been shown to mimic GC infection *in vivo* [13, 54]. Cervical tissue explants were treated with either no hormones, 50 nM estradiol (E2), or a combination of 50 nM E2 and 100 nM progesterone (P4) in phenol red-free media with hormone-stripped fetal bovine serum for 72 h. These concentrations and time frame were chosen to mimic the peaks at the late proliferative phase (E2) and the secretory phase (E2+P4) in the FRT. During the final 24 h, explants were inoculated with piliated Opa-expressing WT MS11 GC at a MOI of 10 GC per luminal epithelial cell. Tissue explants were washed at 6 and 12 h post-inoculation to remove unattached bacteria. Tissues were subsequently cryopreserved, sectioned, stained for GC [55], F-actin, and DNA, and visualized by confocal fluorescence microscopy (CFM) (Fig. 1A). GC colonization was quantified by the percentage of luminal epithelial cells with anti-GC antibody staining (Fig. 1B and 1D) and GC DNA fluorescence intensity (FI) per µm^2^ luminal surface (Fig. 1C and 1E). Hormone treatments, both E2 alone and an E2+P4 combination, significantly increased GC colonization at the endocervix, quantified by both methods (Fig. 1D and 1E). In contrast, the hormone treatments did not significantly impact GC colonization at the ectocervix (Fig. 1B and 1C). Moreover, the menstrual cycle phase of the cervical tissue donors did not significantly affect GC colonization (S1 Table, S1A, S1D, and S1G Fig). Together, these results show that the hormone treatments exclusively increase GC colonization at the endocervix.

**Fig. 1.**
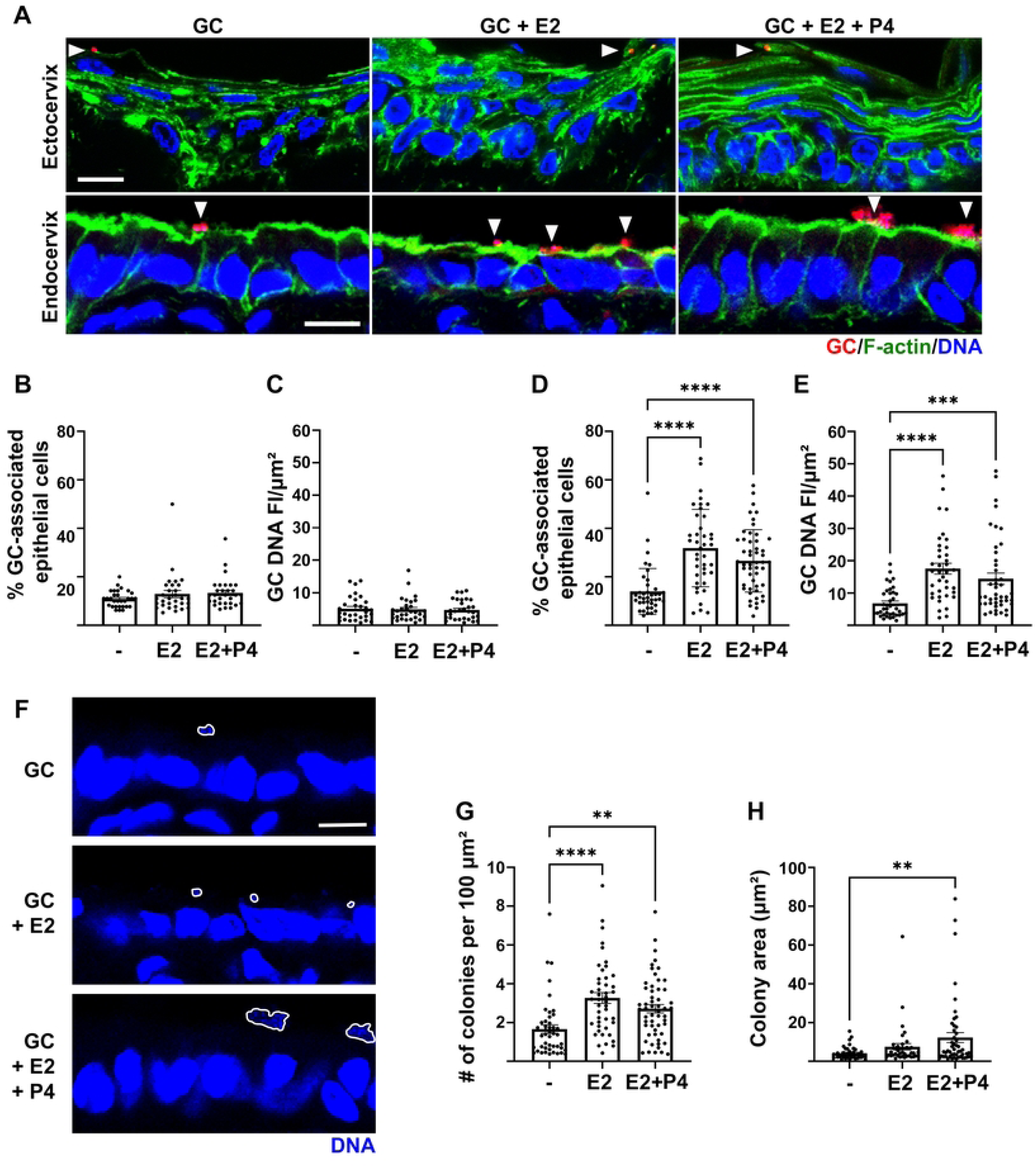
Estradiol and progesterone treatments increase GC colonization exclusively at the endocervix by increasing colony number and size. Human cervical tissue explants were untreated or pretreated for 48 h with estradiol (E2, 50 nM) or a combination of E2 (50 nM) and progesterone (P4, 100 nM). Tissues were incubated with GC (MOI∼10) for 24 h in the absence and presence of the hormones. (A) Representative confocal fluorescence microscopic (CFM) images of the ectocervix (top panels) and endocervix (bottom panels). Arrows indicate GC. Scale bar, 10 µm. (B-E) GC colonization of the ectocervix (B, C) and the endocervix (D, E) was quantified by percentages of luminal epithelial cells associated with GC (B, D) or the fluorescence intensity (FI) of GC DNA staining per μm^2^ at the luminal surface (C, E). (F) Representative CFM images of the endocervix with the areas occupied by colonized GC colonies circled. (G) The number of colonies at the luminal surface (100 µm^2^) of the endocervical epithelium was manually counted. (H) The relative sizes of GC colonies were measured by the area occupied by GC DNA staining of each microcolony. Data points represent individual images. Results are the average values (±SEM) from 5 randomly acquired images per independent analysis, 2 independent analyses per cervix, and cervical tissues from 2-5 donors (>230 epithelial cells total). Data were statistically analyzed using one-way ANOVA. \*\**p*< 0.01, \*\*\**p*<0.001, \*\*\*\**p*<0.0001.

GC can enhance colonization by increasing either the number or the size of microcolonies. We counted the number of GC microcolonies per 100 µm^2^ of the endocervical epithelial luminal surface and measured the areas occupied by individual microcolonies (Fig. 1F-H). Both E2 and E2+P4 treatments resulted in a significantly greater number of colonies on the luminal surface of the endocervix (Fig. 1F and 1G). Further, both hormone treatments increased the average area occupied by GC microcolonies, but only the combination hormone treatment had a significant effect (Fig. 1F and 1H), probably due to the wide range of GC colony sizes. The menstrual cycle phase of cervical tissue donors did not significantly affect the number of GC colonies on the endocervical epithelium (S1B, S1E, and S1H Fig). However, the combination hormone treatment failed to increase the areas occupied by GC microcolonies on the endocervical epithelium of tissue explants from donors at the proliferative phase with internally elevated E2 levels (S1C, S1F, and S1I Fig). Hormone treatments alone also did not significantly alter GC growth (S2 Fig). These results suggest that both E2 alone and in combination with P4 enhance GC colonization at the endocervix, by promoting GC microcolonies attachment to and/or GC-GC aggregation on the luminal surface of the endocervix.

### Hormone treatments do not affect endocervical epithelial cell-cell junctions, epithelial shedding, or GC penetration into the endocervical subepithelium

We previously showed that GC engaging CEACAMs on cervical epithelial cells promotes colonization by inhibiting GC-induced apical junction disruption and epithelial cell shedding, as well as GC penetration into the subepithelium [13]. We examined the impact of hormones on endocervical epithelial apical junction integrity, epithelial shedding, and GC penetration. Junction integrity was evaluated by determining the ratio of E-cadherin FI at the cell-cell junction relative to that at the adjacent cytoplasm using E-cadherin FI line profiles (Fig. 2A). The E-cadherin junction to cytoplasmic FI ratio (FIR) was not significantly changed in the presence of hormones compared to no hormone controls (Fig. 2B). Endocervical epithelial shedding was evaluated by counting the percentage of epithelial cells disassociating from the epithelium (Fig. 2C, arrow), but no differences were found between no hormone and hormone treatments (Fig. 2D). GC penetration was quantified by the percentage of GC-associated cells with GC at the basolateral membrane of epithelial cells and at the subepithelium (Fig. 2E and 2F) and the percentage of total GC FI below the basal surface of the epithelium (Fig. 2G). No changes in GC penetration levels were detected by either method, regardless of hormone treatment (Fig. 2E-G). However, it was noted that the E2-only treatment, but not the E2+P4 combination, altered the shape of endocervical epithelial cells from columnar to cuboidal (Fig. 2H). This shape change was quantified by the ratio of cell height to width of individual endocervical epithelial cells (Fig. 2I) and the number of cells per 100 µm^2^ luminal surface (Fig. 2J). E2 but not the E2+P4 combination significantly decreased the height-to-width ratio (Fig. 2I) and the number of endocervical epithelial cells per 100 µm^2^ luminal surface (Fig. 2J). These results show that hormone treatments do not significantly interrupt the apical junction of the endocervical epithelial cells or increase epithelial shedding and GC penetration, but the E2 treatment changes the shape of endocervical epithelial cells.

**Fig. 2.**
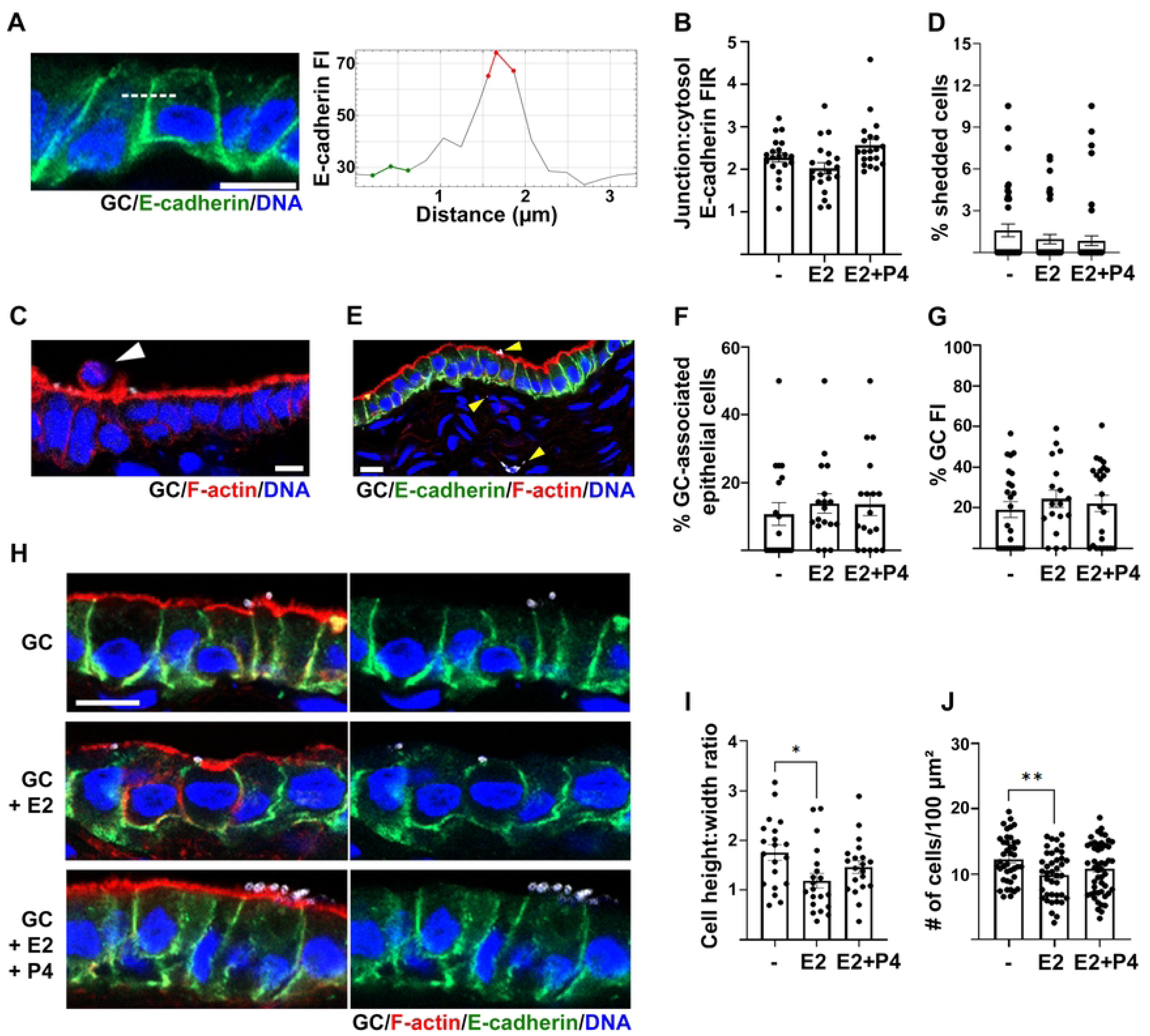
Hormone treatments have no effect on the endocervical epithelial cell-cell junction and shedding or GC penetration into the endocervical subepithelium. Human cervical tissue explants were treated and stained as described in Fig. 1. **(A, B)** The integrity of endocervical epithelial cell-cell junctions was quantified by ratios of E-cadherin FI at the cell-cell junction to the cytoplasm (**B**) using FI line profiles across the cell-cell junction **(A**). (**C**) A representative CFM image shows endocervical epithelial cell shedding (arrow). (**D**) Endocervical epithelial shedding was quantified by the percentage of epithelial cells moving out of the monolayer. (**E**) A representative CFM image shows penetrated GC in the endocervix (arrows). (**F, G**) GC penetration was quantified by the percentage of GC-associated cells with GC staining at the basal surface or subepithelium (**F**) and the percentage of GC fluorescence in the subepithelium (**G**). (**H**) Representative images show the shape changes of endocervical epithelial cells by hormone treatments. (**I, J**) The shape of endocervical epithelial cells was evaluated by the ratio of cell height to width (**I**) and the number of epithelial cells per µm^2^ (**J**). Scale bar, 10 µm. Data points represent individual images. Results are the average values (±SEM) from 5 randomly acquired images, 2 independent analyses per cervix (>100 epithelial cells total), and 2-5 cervixes. Data were statistically analyzed by Student’s *t*-test. \**p*<0.05; \*\**p*< 0.01.

### Hormone-induced increases in GC colonization at the endocervix are not the result of increased CEACAM expression or GC-hormone interactions

The interactions of GC surface molecules, Opas, with their human-specific receptors, CEACAMs, increase GC colonization at the endocervix and ectocervix of tissue explants [13]. We hypothesized that sex hormones facilitate GC colonization by upregulating CEACAM expression at the luminal surface of the endocervix. To test this hypothesis, we analyzed the expression level and cellular distribution of CEACAMs in the endocervical epithelial cells in response to hormone treatment using immunofluorescence microscopy (IFM). The expression level of CEACAMs in endocervical epithelial cells was evaluated by the FI of CEACAM staining per cell (Fig. 3A-C) and the luminal surface expression of CEACAMs by calculating the percentage of total CEACAM FI (Fig. 3A, 3D, and 3E) at the apical surface of endocervical epithelial cells. Hormone treatments did not significantly change CEACAM FI per epithelial cell (Fig. 3C). However, treatment with E2, but not E2+P4, significantly reduced the percentage of CEACAMs at the apical surface of endocervical epithelial cells (Fig. 3A, 3D, and 3E). IFM images showed that E2 treatment induced a redistribution of CEACAMs from the apical surface to intracellular puncta (Fig. 3A, middle panels). Notably, CEACAM redistribution was not observed in tissue explants treated with hormones alone without GC inoculation (S3 Fig), indicating a synergistic effect of E2 and the GC. These data suggest that hormone treatments do not increase CEACAM expression at the apical surface of endocervical epithelial cells, and therefore, CEACAMs do not contribute to hormone-enhanced GC colonization.

**Fig. 3.**
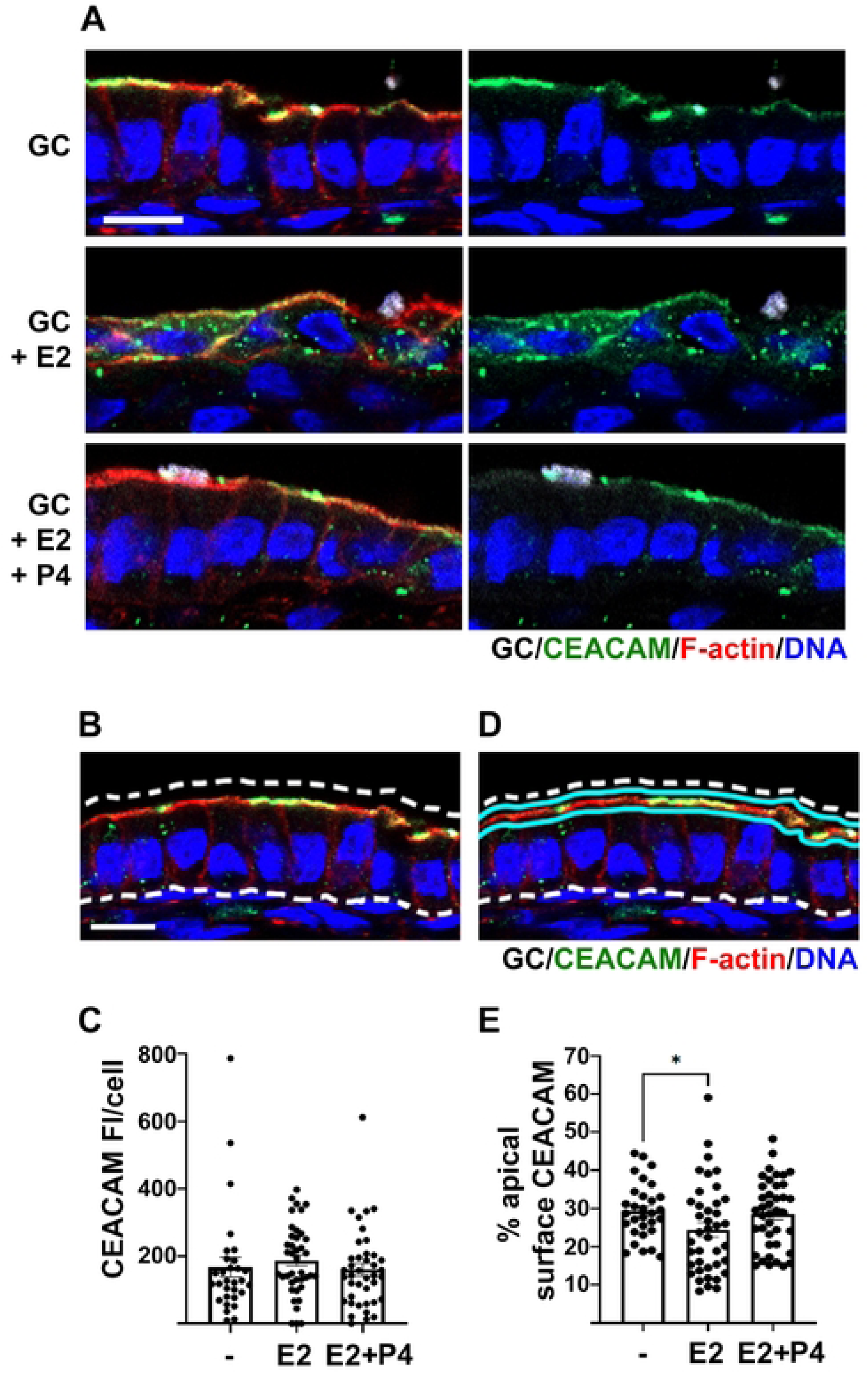
Hormone-induced increase in GC colonization on the endocervix is independent of CEACAMs. Cervical tissue explants were treated with hormones and inoculated with MS11 WT GC as described in Fig. 1. Tissue sections were stained for CEACAMs (1, 3, and 6), F-actin, and DNA. (**A)** Representative images of the endocervical epithelium. (**B, C**) The expression level of CEACAMs was evaluated by measuring CEACAM FI per epithelial cell. (**D, E)** The relative expression level of CEACAMs at the luminal surface of endocervical epithelial cells was evaluated by the percent of CEACAM FI on the apical surface. Scale bar, 10 µm. Data points represent individual images. Results are the average values (±SEM) from 5 randomly acquired images per analysis, 2 independent analyses per cervix (>230 epithelial cells total), and cervixes from 3-4 donors. Results were statistically analyzed using Student’s *t*-test. \**p*<0.05.

P4 has been shown to enhance *E. coli* self-aggregation and promote *E. coli* and *B. fragilis* biofilm formation [56]. To determine if hormones directly impact GC aggregation, we incubated GC, which had been vigorously vortexed to break aggregates, with hormones for 6 h. The concentrations of hormones used here were the same as those for treating tissue explants described above. Bacteria were stained with the DNA dye SYTO9, and Z-stack images were acquired. Z-stacks of images with GC aggregates on top of each other were excluded. The occupied areas and mean fluorescence intensity (MFI) of individual GC aggregates were measured using the sum intensity projections of images (Fig. 4A). Hormone treatments did not significantly change the occupied area (Fig. 4B) or the MFI of GC aggregates (Fig. 4C). These results suggest that E2 and P4 do not directly impact GC aggregation.

**Fig. 4.**
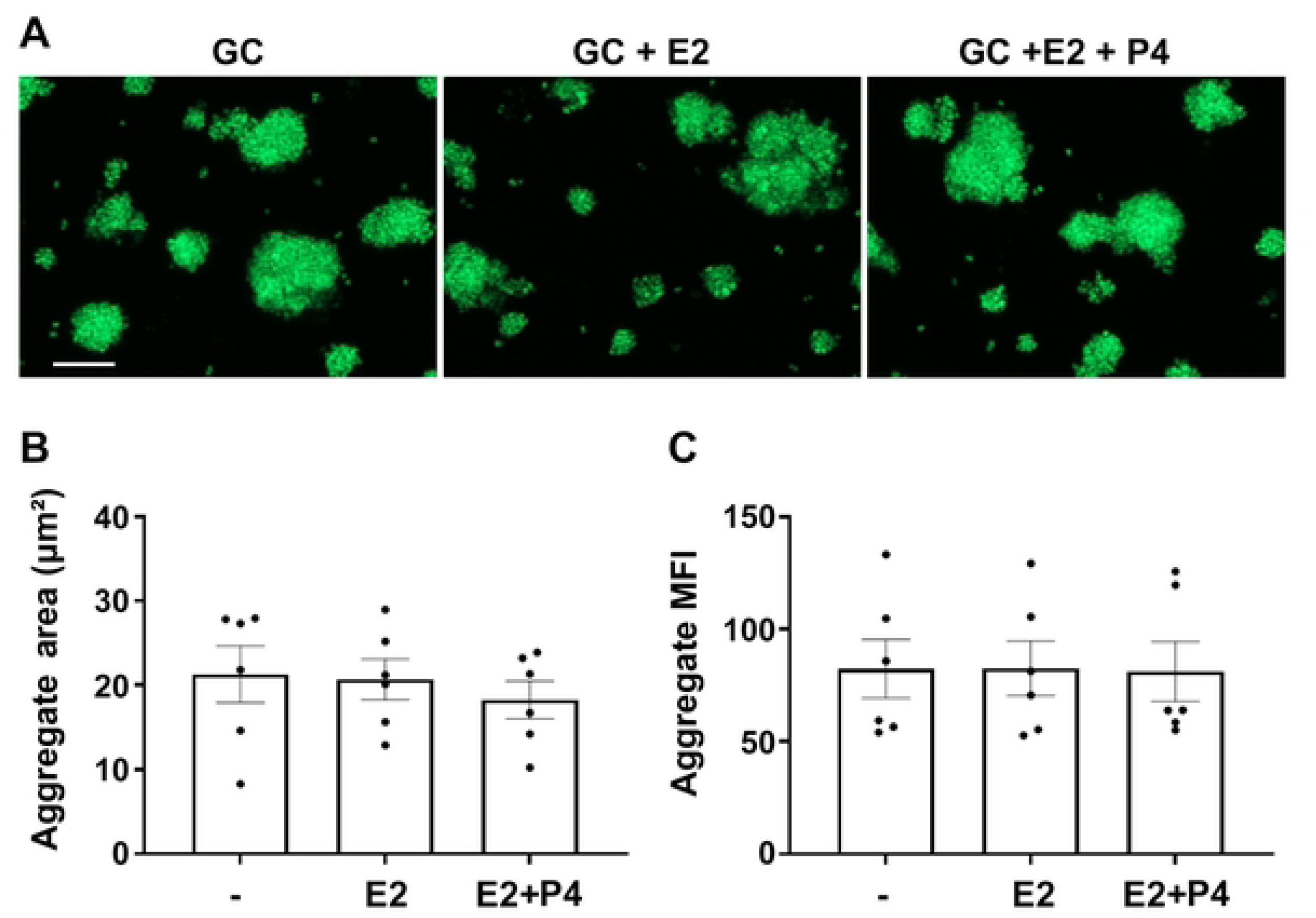
Sex hormones do not directly regulate GC-GC aggregation. Piliated WT MS11 GC were incubated with 50 nM E2 or 50 nM E2 and 100 nM P4 in coverslip chambers at 37°C and 5% CO_2_ for 6 h, stained with the DNA dye SYTO9. Z-stack images were randomly acquired using CFM. (**A**) Representative images with sum intensity projections are shown. Scale bar, 10 µm. (**B**) The average size of bacterial aggregates was evaluated by the area each aggregate occupied. (**C**) The bacterial densities of individual aggregates were measured by the mean fluorescence intensity (MFI) of SYTO9 staining. Shown are average values (±SEM) from 3 independent experiments, 2 wells per experiment, >3 randomly acquired images per well. Each data point represents one well. Results were statistically analyzed using two-way ANOVA.

### GC microcolonies reduce the membrane-anchored mucin protein MUC1 at adherent sites of the endocervix

Mucus is a potential factor influencing GC colonization since its production by the endocervical epithelial cells varies with the hormonal cycles [39, 40, 50, 51]. Mucus, consisting of membrane-anchored and gel-forming mucin layers, provides a protective barrier against microbial attachment to host cells [41, 42]. To examine whether hormone treatments affect GC interactions with the endocervical mucus during colonization, cervical tissue explants treated with and without hormones and inoculated with GC were stained for the membrane-anchored mucin MUC1, GC, and DNA (Fig. 5A). The total MUC1 FI per endocervical epithelial cell was measured as an indication of relative MUC1 protein expression level (Fig. 5B), and MUC1 FI per μm^2^ of the luminal surface was measured as an indication of MUC1 luminal surface expression level (Fig. 5C). Neither MUC1 FI per cell nor per μm^2^ of the luminal surface were affected by hormone treatments (Fig. 5A-C), suggesting that MUC1 expression levels do not vary in response to the hormone treatments, consistent with previous reports that MUC1 levels are maintained throughout the menstrual cycle [39, 57]. Interestingly, MUC1 staining was noticeably reduced under GC microcolonies in both hormone-treated and non-treated tissue explants (Fig. 5D). This reduction was quantified by the ratio of MUC1 FI under GC microcolonies relative to MUC1 FI in an adjacent uncolonized location of equivalent size on the luminal surface of the endocervix. The average MUC1 FI under GC microcolonies was lower than that at areas without GC colonization under all the hormone treatment conditions, leading to the FI ratios smaller than 1 (Fig. 5E). These results suggest that GC can diffuse through the mucus barrier to interact with the epithelial cell surface despite hormone treatment.

**Fig. 5.**
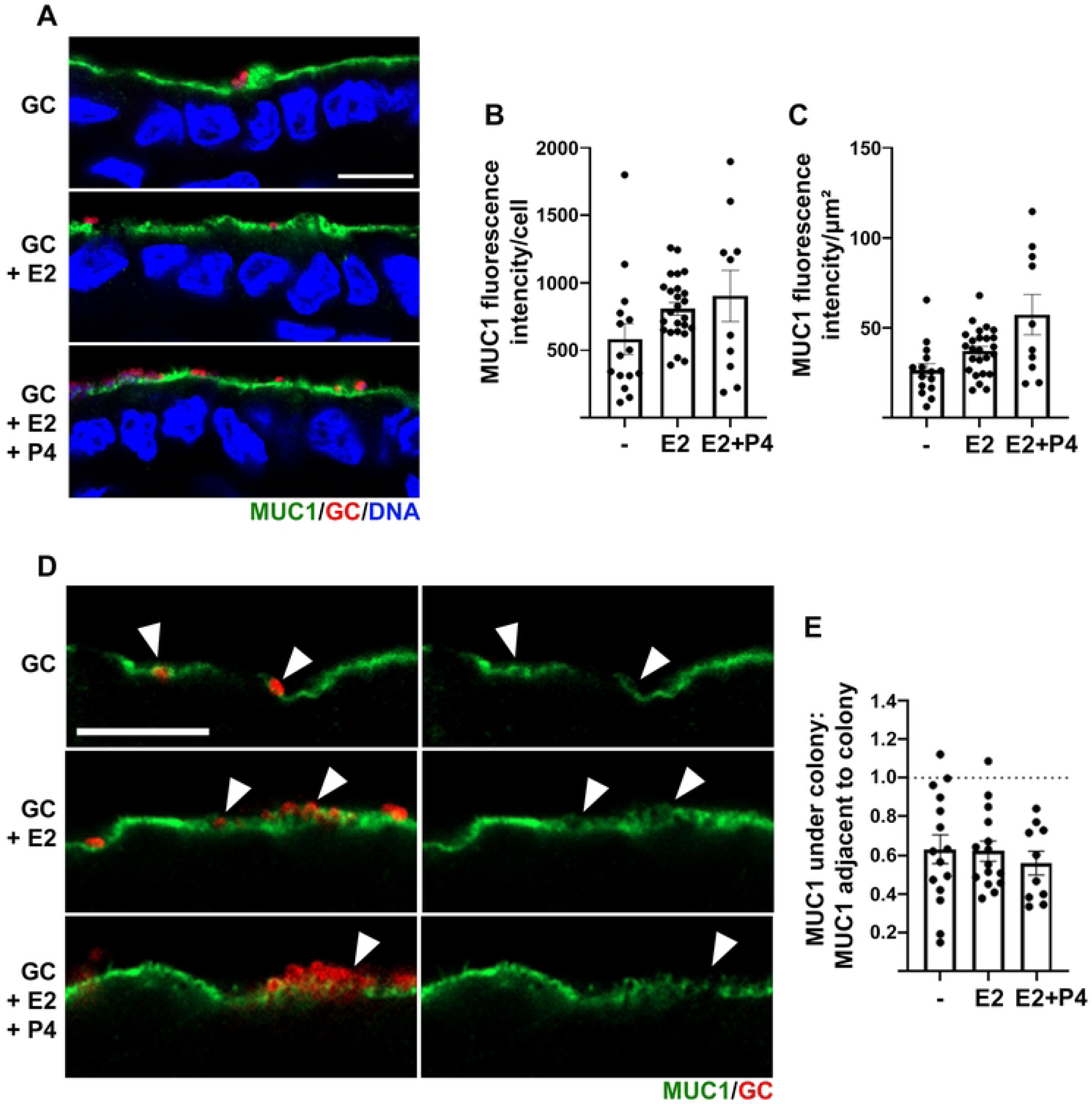
GC microcolonies reduce the membrane-anchored MUC1 at adherent sites of the endocervical epithelium. Cervical tissue explants were treated with hormones and incubated with GC, as described in Fig. 1. Tissue sections were stained for MUC1, GC, and DNA. (**A**) Representative CFM images. (**B**) The expression level of MUC1 was estimated by the FI of MUC1 per epithelial cell. (**C**) The luminal surface expression level of MUC1 was evaluated by the FI of MUC1 along the luminal surface. (**D**) Representative CFM images show MUC1 staining at GC colonizing sites of the luminal surface (arrows). (**E**) The ratio of MUC1 FI under GC colonies relative to the MUC1 FI in an adjacent area without GC. Scale bar, 10 µm. Data points represent individual images. Results are the average values (±SEM) from 5 randomly acquired images per cervix (>100 epithelial cells total) and cervixes from 2-3 donors. Data were statistically analyzed using Student’s *t*-test.

### GC can effectively diffuse through MUC5AC-dominated gels, and MUC5B slows GC diffusion

Gel-forming mucins, including MUC5AC and MUC5B, form a mucus gel layer and prevent contact of microbes with the membrane-anchored mucins [42, 45]. E2-induced MUC5B production at the cervix reduces the viscoelasticity of the cervical mucus, facilitating sperm migration [40, 43, 51]. To determine if sex hormone-induced changes in mucin composition impact GC diffusion through the gel-forming mucin layer, we generated hydrogels using commercially available animal mucins [58, 59]. MUC5B and MUC5AC mucins were mixed in various ratios to a final concentration of 2% (w/v) and crosslinked via disulfide bonding using PEG-4SH to create hydrogels with ∼1 mm thickness on coverslips. Rheological analysis showed decreases in hydrogel viscoelasticity with increasing MUC5B concentration (S4 Fig). GC labeled with the DNA dye SYTO9 were added to the top of the hydrogels and incubated for 6 h at 37°C. Z-stack images were acquired from the bottom of the hydrogels up to 35 μm (Fig. 6A). GC diffusion efficiency in the mucin hydrogel was estimated by the GC MFI of each Z-slice over the distance from the coverslip (Fig. 6B) and the percentage of GC FI of each Z-slice relative to the total GC MFI in the all Z-slices (Fig. 6C). GC were detected in the bottom of hydrogels with different MUC5AC and MUC5B ratios (Fig. 6A), showing that GC, added from the top of hydrogels, reached coverslips. GC MFI and FI% all peaked close to the coverslip and gradually decreased as the distance from the coverslip increased (Fig. 6A-C). To evaluate the interference of hydrogels with CFM imaging, hydrogels incubated with a fluorescent antibody for 6 h were analyzed using CFM. Z-stack images showed that antibody-associated fluorescence decreased at a linear rate as the focal plane moved up into the hydrogel (Fig. 6D). Together, these data indicate that GC reached the bottom of all hydrogels tested. However, GC MFI and FI% decreased in image slices close to the coverslip (4 µm) but increased in image slices away from the coverslip (10-25 µm) as MUC5B concentration was increased (Fig. 6A-C). Hydrogels containing only MUC5B exhibited higher levels of background fluorescence and thus were excluded from analysis. These results suggest that while GC can successfully penetrate through gel-forming mucus, GC diffusion is less efficient in MCU5B-dominated than MUC5AC-dominant mucus.

**Fig. 6.**
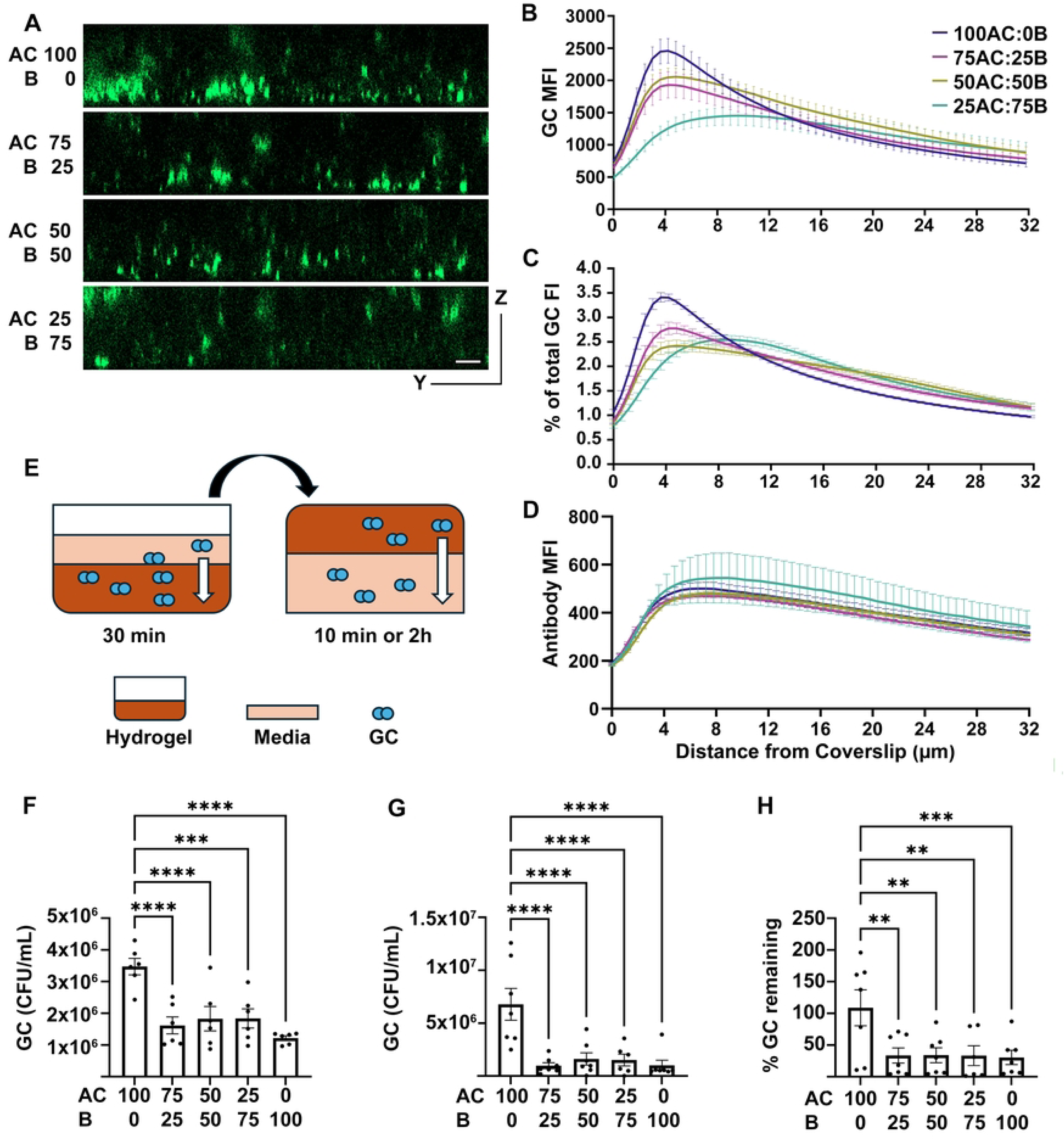
GC penetrate MUC5AC-dominant mucin hydrogels more efficiently than MUC5B-dominant mucin hydrogels *in vitro*. Mucin hydrogels (2%, ∼1 mm height) containing various fractions of porcine MUC5AC and bovine MUC5B were polymerized on cover glasses by 2% PEG-SH. Piliated WT MS11 GC (5×10^3^) stained with SYTO9 were added to the top of the hydrogel and incubated for 6 h. Z-stack images were acquired from the cover glass up by CFM. **(A)** Representative yz images of hydrogels from cover glass (bottom) up to 32 µm. Scale bar, 10 µm. **(B, C)** GC MFI (±SEM) of individual slices **(B)** and percentages of total Z-stack FI **(C)** were plotted over ascending distance from the coverslip. **(D)** Alexa Fluor 488-goat anti-chicken IgG (20 µg in 10 µL) was added to the top of hydrogels, incubated for 6 h at 37°C, and imaged as described above. The MFI of antibodies was plotted versus the ascending distance from the coverslip. **(E-H)** Mucin hydrogels with various MUC5AC and MUC5B concentrations in 96-well plates were incubated with MS11 GC (1.28×10^7^ GC in 30 µL) from the top for 30 min at 37°C and 5% CO_2_. Then, wells were filled with media, sealed, flipped upside down, and incubated for either 10 min or 2 h **(E)**. After the media were drained, GC were released from the hydrogels and enumerated. Shown are the average CFU (±SEM) remaining in the hydrogels after 10 min **(F)** or 2 h **(G)** upside-down incubation and the average percentages of GC remaining in the hydrogels after 2 h upside-down incubation relative to those after 10 min upside-down incubation **(H)**. The results were generated from 3 independent experiments. Data points represent individual hydrogels **(F-H)**. Data were statistically analyzed by two-way ANOVA. \*\**p*< 0.01; \*\*\**p*<0.005; \*\*\*\**p*<0.0001.

To investigate why GC diffusion is less efficient in relatively low viscoelastic MUC5B-dominan versus relatively high viscoelastic MUC5AC-dominated hydrogels, we inoculated hydrogels with different MUC5B:MUC5AC ratios from the top with GC and allowed GC to diffuse into hydrogels for 30 min rather than 6 h. After 30 min of incubation, wells were filled with media, sealed, and then turned upside down for 10 min and 2 h (Fig. 6E). Bacteria remaining in the hydrogels were quantified. The 10-min upside down incubation allowed GC that had not entered hydrogels to move away from hydrogels by gravity, and therefore, the data reflect the number of GC diffused into hydrogens during the 30-min bacterial loading (Fig. 6F). The 2-h upside down incubation allowed GC that entered hydrogels during the 30-min loading to disassociate from the hydrogels by gravity, and therefore, the data quantified the number of GC that remained in hydrogels against gravity (Fig. 6G). Notably, the number of GC diffused into 100% MUC5AC hydrogels during the 30-min loading (10 min upside-down incubation) (Fig. 6F) and the number of GC remained in 100% MUC5AC hydrogels after the 2-h upside-down incubation (Fig. 6G) were all significantly higher than in any hydrogels containing MUC5B. Further, much higher percentages of GC that diffused into 100% MUC5AC hydrogels during the 30 min loading remained in the hydrogels after the 2 h upside-down incubation than in any hydrogels containing MUC5B (Fig. 6H). These data further confirm that GC diffuse through MUC5AC hydrogels more efficiently than through hydrogels containing MUC5B and also suggest that GC can remain in MUC5AC but not MUC5B hydrogels against gravity.

### Cervical mucus promotes GC aggregation

To test whether hormone-induced mucus alterations contribute to GC colonization, we concentrated mucus from the supernatants of cervical tissue explants treated with or without hormones and infected with GC using Amicon Ultra centrifugal filters (100 kDa) and quantified their mucin concentrations. Western blot analysis detected MUC5B in cervical explant supernatants from most donors, while MUC5AC was detected in some (S5 Fig). Well-dispersed GC were incubated with cervical mucus (0.1% mucins w/v) from explant supernatants in coverslip chambers for 6 h, stained with SYTO9, and imaged by CFM. GC aggregation was quantified by measuring the areas occupied by individual GC aggregates, using a 2D sum stack of Z-stacks obtained by CFM. Cervical mucus, regardless of whether it came from hormone-treated or non-treated tissue explants, significantly increased the sizes of GC aggregates compared to medium controls (Fig. 7A and 7B). Furthermore, the average aggregate sizes of GC incubated with cervical mucus from E2-treated tissue explants were significantly smaller than GC incubated with cervical mucus from tissue explants not treated with hormones (Fig. 7B). However, differences between the aggregate size of GC incubated with cervical mucus from E2 plus P4-treated and hormone-free tissues were not statistically significant (Fig. 7B). Further analysis showed that cervical mucus generated a wide range of large GC aggregates (>300 µm^2^) that were not detected in the absence of cervical mucus (Fig. 7D and 7E). These results indicate that the mucus secreted by the cervix promotes GC-GC aggregation.

**Fig. 7.**
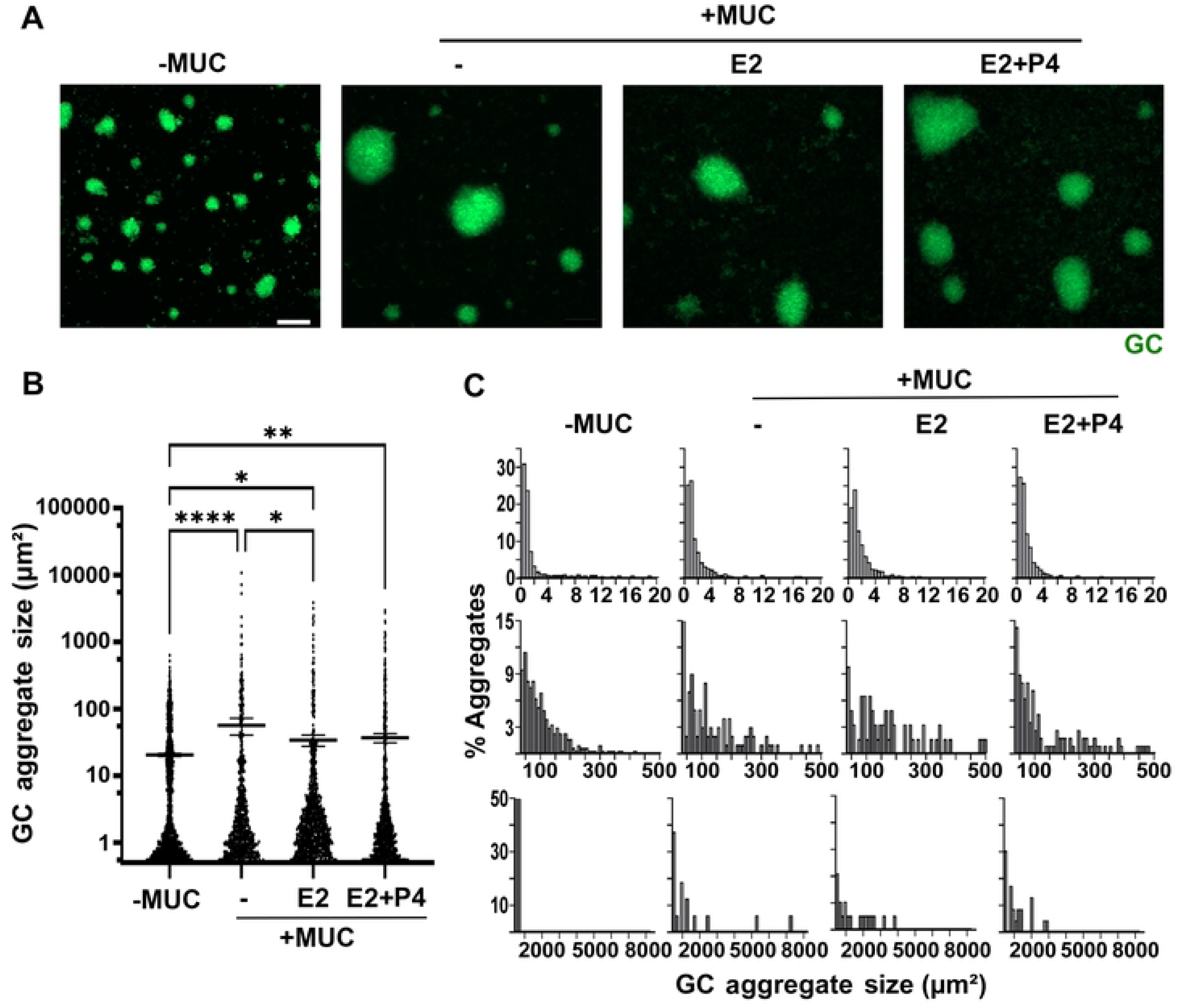
Cervical mucus promotes GC aggregation. Post-inoculation supernatants were collected, and mucins were concentrated using molecular filters with a 100 kDa cutoff. Piliated MS11 GC were incubated with the mucus (0.1%) for 6 h at 37°C and 5% CO_2_, stained with the DNA dye SYTO9, and imaged by CFM. (**A)** Representative maximal projections of Z-stack CFM images. Scale bar, 20 µm. (**B**) Sizes of individual aggregates were measured by their occupied area using maximal projection images. (**C**) Percentage of GC aggregates at 1-20 (the first row), 21-500 (the second row), and 501-8000 (the third row) µm^2^ sizes. Data were generated from 2 independent experiments using cervical mucus of 4 donors and 6 randomly acquired Z-stack images per condition per experiment. Results were statistically analyzed by two-way ANOVA. \**p*<0.05; \*\**p*< 0.01; \*\*\*\**p*<0.0001.

### MUC5AC promotes GC aggregation more efficiently than MUC5B

The MUC5B level at the cervix is modulated by the levels of E2, but the MUC5AC level is not changed by the menstrual cycle [43, 50]. To determine how the MUC5B and MUC5AC composition of the cervical mucus impacts GC aggregation, well-dispersed GC were incubated with 2% (w/v) animal mucins with various ratios of MUC5B to MUC5AC for 6 h before SYTO9 staining and imaging. Animal mucins used here were not crosslinked via disulfide bonds, because the crosslinking interfered with CFM analysis. GC aggregation was evaluated by determining the area individual aggregates occupied using a 2D sum stack of Z-stacks obtained by CFM. Similar to mucus collected from cervical tissue explant culture supernatants, uncrosslinked animal mucins of all tested MUC5AC to MUC5B ratios increased the average sizes of GC aggregates, compared to no mucin controls (Fig. 8A and 8B). However, GC aggregates reduced their average sizes as the proportion of MUC5B increased (Fig. 8B). Animal mucins enabled GC to aggregate into sizes larger than 300 µm^2^ while MUC5B primarily reduced the number of very large aggregates (>1000 µm^2^) (Fig. 8C). Under the same conditions, there was no significantly increase in the number of GC during the 6-h incubation with animal mucins (S6 Fig), suggesting that enhanced GC aggregation is not the result of increased growth. These results further confirm the enhancement effect of mucus on GC-GC aggregation.

**Fig. 8.**
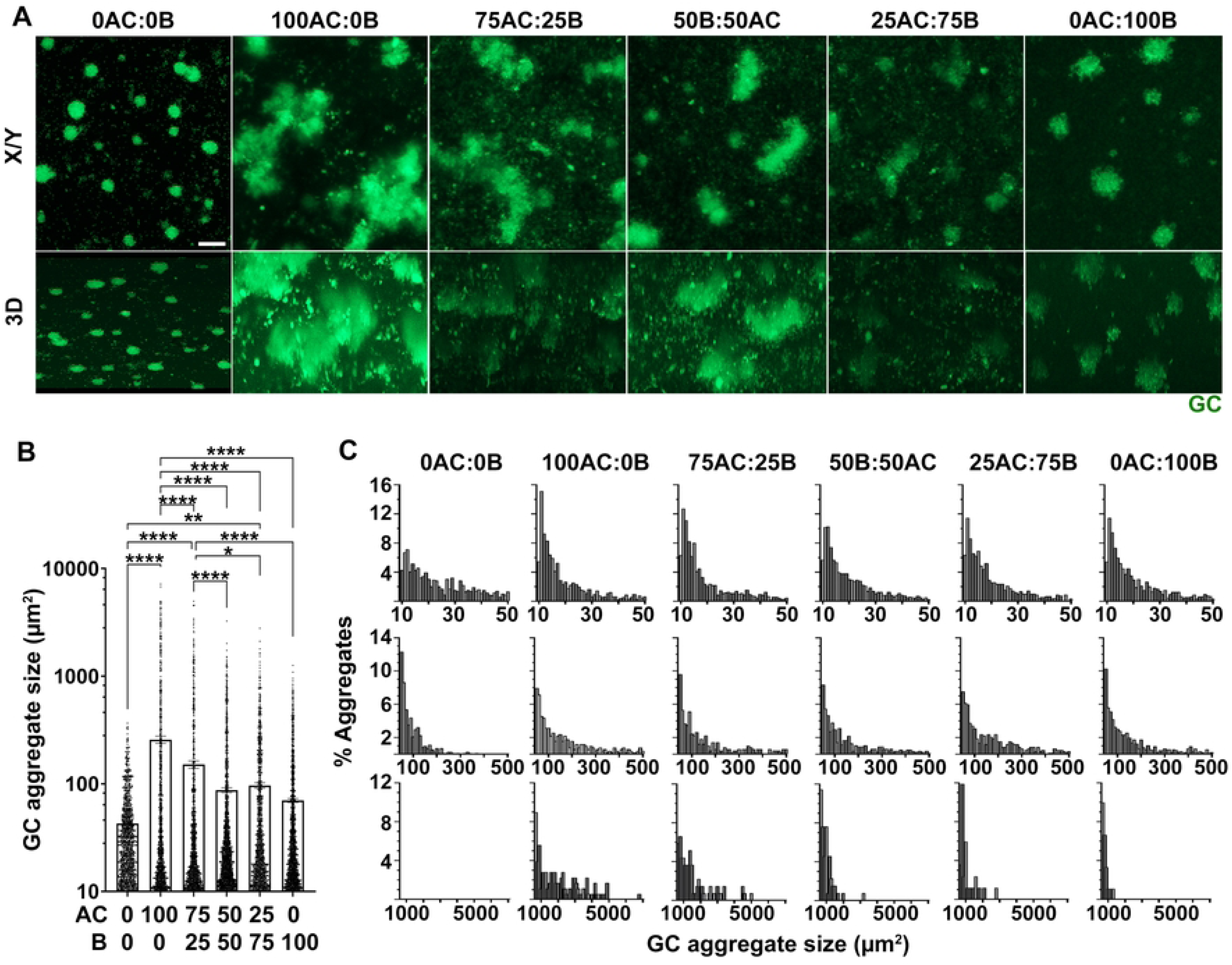
MUC5AC promotes GC aggregation more efficiently than MUC5B. Piliated MS11 GC were incubated with a 2% mucin protein mixture of varying MUC5B:MUC5AC ratios on coverslips for 6 h, stained with the DNA dye SYTO9, and imaged by CFM. (**A**) The XY view of representative CFM sum stacks (top panels) and the 3D projection of representative Z-stack images (bottom panels). Scale bar, 20 µm. (**B**) Average sizes (±SEM) of individual aggregates were measured by occupied areas. (**C**) Percentages of GC aggregates at 1-50 (the first row), 51-500 (the second row), and 501-8000 (the third row) µm^2^ sizes. Data were generated from 3 independent experiments and >3 randomly acquired Z-stack images per condition per experiment. \**p*<0.05; \*\**p*< 0.01; \*\*\*\**p*<0.0001. Results were statistically analyzed by two-way ANOVA.

To determine the effect of E2-induced increases in MUC5B production on GC-GC aggregation during the proliferative stage of the menstrual cycle, we incrementally increased animal MUC5B in the uncrosslinked mucin mixture from 0.5 to 2% (w/v) while keeping MUC5AC at 0.5%. This changed the MUC5AC to MUC5B ratio from 50:50 to 10:80 and increased the total amount of mucins from 1 to 2.5% (w/v). Again, animal mucins with different amounts of MUC5B and the same amount of MUC5AC all enhanced GC-GC aggregation, compared to no mucin control (Fig. 9A and 9B). However, the sizes of GC aggregates decreased as the amount of MUC5B increased in the uncrosslinked mucin mixture (Fig. 9A and 9B), with a drastic decrease in the number of GC aggregates larger than 300 μm^2^ (Fig. 9C). Our data suggest that the aggregation enhancement effect can be modulated by mucin composition changes over the hormonal cycle, with MUC5AC-dominant mucus (at low hormone or high progesterone phases) promoting GC-GC aggregation more effectively than MUC5B-dominant mucus (at the high E2 phase).

**Fig. 9.**
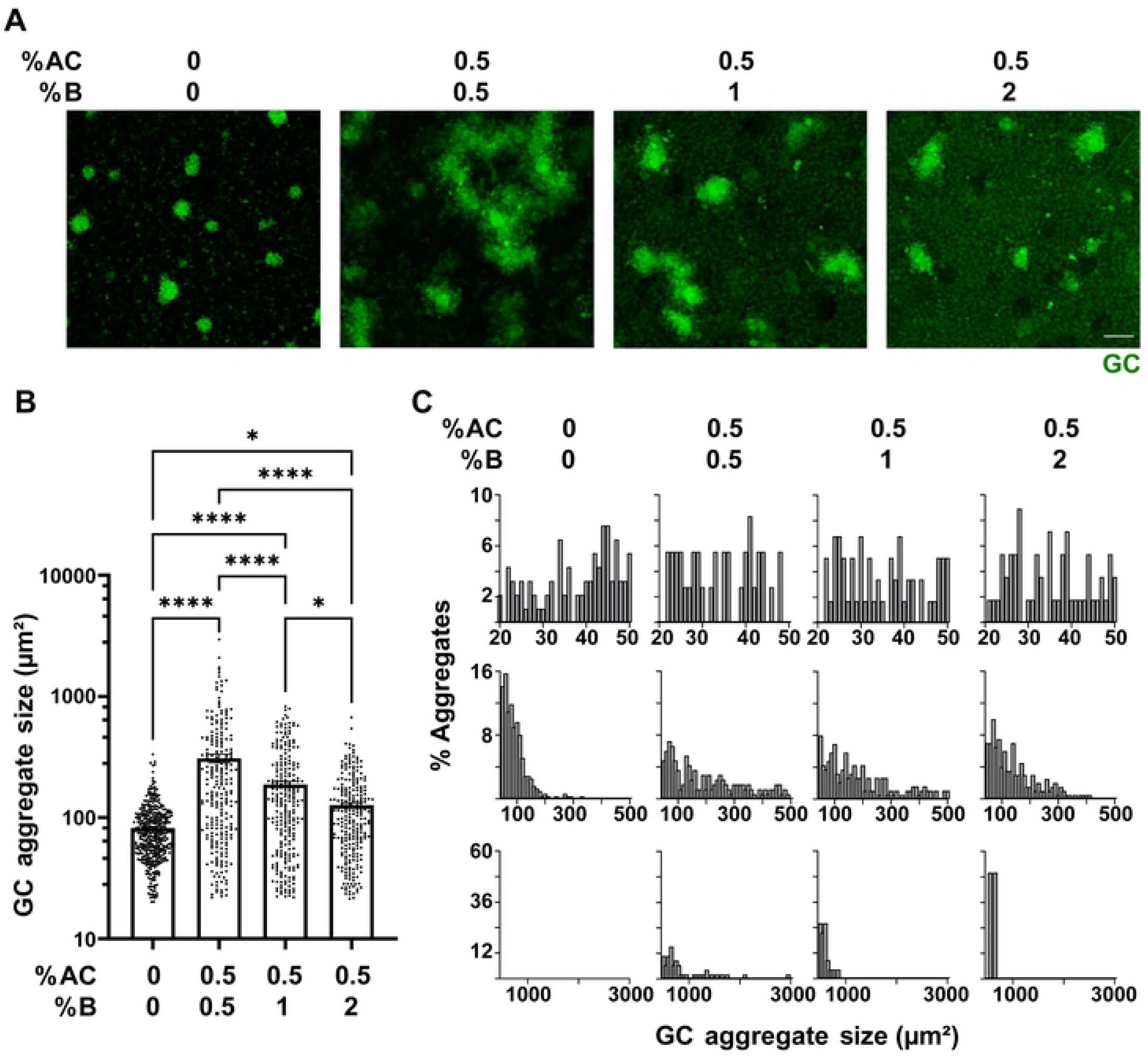
MUC5B reduces the ability of mucins to enhance GC aggregation. Piliated MS11 GC were incubated with a mucin mixture containing 0.5% MUC5AC mixed with 0.5, 1, and 2% MUC5B on coverslips for 6 h, stained with the DNA dye SYTO9, and imaged by CFM. (**A**) The XY view of representative CFM sum stacks. Scale bar, 20 µm. (**B**) Average sizes (±SEM) of individual aggregates were measured by occupied areas. (**C**) Percentages of GC aggregates at 1-50 (the first row), 51-500 (the second row), and 501-8000 (the third row) µm^2^ sizes. Data were generated from 3 independent experiments and 10 randomly acquired Z-stack images per condition per experiment. \**p*<0.05; \*\*\*\**p*<0.0001. Results were statistically analyzed by two-way ANOVA.

## Discussion

Using the human cervical tissue explant model, which can mimic GC infection *in vivo*, this study demonstrates that a 72-h treatment with E2 alone or in combination with P4 increases MS11 WT GC colonization exclusively at the endocervix without significantly altering the level of GC transmigration across the epithelium. The hormone treatments enhanced GC colonization by increasing the number of GC microcolonies at the luminal surface of the endocervical epithelium, and the E2+P4 combination also increased the size of GC microcolonies, a consequence of enhanced GC aggregation. However, the hormone treatments did not increase the expression of the GC host receptor CEACAMs, reduce endocervical epithelial cell shedding, or directly promote GC aggregation in the absence of cervical tissue explants, eliminating three possible mechanisms underlying hormone-enhanced GC colonization. Instead, mucus from the supernatants of cervical tissue explants treated with or without hormones, as well as animal mucin protein mixtures with compositions mimicking the gel-forming mucins at different stages of the hormone cycle, enhanced GC-GC aggregation. Furthermore, GC can effectively penetrate the protective mucus layers on the luminal surface of hormone-treated and untreated cervical tissue explants in *ex vivo* conditions, as well as mucin hydrogels *in vitro*, converting the endocervical mucus layer from a colonization barrier to a colonization facilitator.

Previous studies [9, 35, 60], including our data [61], showed that E2 can weaken the epithelial cell-cell junctions of primary cervical and uterine epithelial cells and cell lines and increase epithelial paracellular permeability *in vitro*. Loss of epithelial cell junction integrity has been shown to promote epithelial shedding, a mechanism by which the host eliminates bacteria colonizing epithelial cells [20, 62, 63]. GC-induced apical junction disruption also facilitates bacterial penetration into the subepithelium [13, 20]. Thus, we initially hypothesized that sex hormone treatments would reduce GC colonization and increase GC penetration by compromising the cervical epithelium. However, our findings indicate that sex hormones promote GC colonization at the endocervix without affecting the apical junction integrity or epithelial shedding (Fig. 2), contradicting previously published data. While the reasons underlying these differences in results are unclear, there are many differences between our cervical tissue explant *ex vivo* model and the *in vitro* epithelial cell culture models, among which two stand out. First, the type and/or levels of specific mucins produced by epithelial cells from different regions of the FRT and in different model systems likely differ. This study shows that mucus from tissue explant culture media and uncross-linked animal mucin protein mixtures, with compositions similar to those of cervical gel-forming mucins, promote bacterium-bacterium aggregation, a mechanism that facilitates GC attachment [64, 65]. Furthermore, enhanced GC colonization was only observed in the endocervix (Fig. 1), where columnar epithelial cells are the primary producers of cervical mucins [39, 40], underscoring the role of cervical-generated mucus in hormone-enhanced GC colonization. Second, the local cytokine environment of the tissue explant model is likely distinct from the epithelial cell culture models, as human cervical tissue explants preserve the heterogeneous epithelium and subepithelial and tissue immune cells [66–68]. Our recently published data [69] show that human cervical tissue explants constitutively secrete the anti-inflammatory cytokine IL-10, and GC inoculation further increases the level of IL-10 secretion. Importantly, IL-10 protects the cervical epithelia from damage by inflammatory cytokines and inhibits GC-associated cervical epithelial cells from shedding [69]. The epithelial protective function of locally secreted IL-10 potentially prevents E2 from weakening epithelial cell-cell junctions. Sex hormones can also regulate the local production of inflammatory cytokines [70, 71]. While additional contributing factors from human cervical tissue explants remain to be discovered, hormone-induced mucin and IL-10 production potentially inhibits hormone-induced epithelial junction disruption and enhance GC colonization at the endocervix.

A significant finding of this study is the role of mucus in facilitating rather than preventing GC colonization at the endocervix, revealing a novel mechanism by which GC can evade host defense. GC can not only effectively migrate through the mucus (Figs. 5 and 6), but also increase self-aggregation in the presence of mucus or mucin protein mixtures (Figs. 7-9). Mucins constitute a family of large, heavily glycosylated proteins that represent the primary structural components of the mucus barrier, including membrane-anchored mucins, such as MUC1 and MUC4, and secreted or gel-like mucins, such as MUC5AC and MUC5B at the cervix [41, 45]. Besides lubricating the mucosal surface, the mucin layers serve as a physical barrier, preventing pathogens from directly interacting with epithelial cells by blocking microbial diffusion towards the epithelial cells and entrapping pathogens within the mucus. Mucus also contains secreted anti-microbial reagents, including antibodies, peptides, glycans, and complement factors that can neutralize entrapped pathogens [72, 73]. The mechanism by which GC evade complement-mediated killing has been well studied [74]. However, the mechanism by which GC overcome the mucus barrier to aggregate on the cervical epithelium is unknown. As both mucins and the GC surface are highly negatively charged, electrostatic repulsion between GC and mucins should prevent GC from getting close to mucus [75, 76]. The type IV pili, which have positively charged amino acid patches at the PilE tip [77, 78], may help GC attach to and migrate through the mucus layer. Pili are required for GC to attach to each other [12, 14] and endocervical epithelial cells [13], whose surface is also generally negatively charged. In addition, other GC surface molecules, including opacity-associated proteins (Opa) and lipooligosaccharides (LOS), are responsible for intimate GC-GC and GC-host cell interactions, after the initial pili contraction brings GC close to each other or to epithelial cells and their host receptors [15, 18, 79]. The increased GC aggregation in the presence of mucus or mucins indicates a preference of GC to interact with each other over mucus. This preference suggests that pili-mucus interactions are relatively transient and weak, insufficient to completely overcome the electrostatic repulsion of negative charges, preventing GC from tightly attaching to mucus for entrapment. Nonetheless, how GC overcome the negatively charged mucus barrier remains an open question.

Cervical mucus, as a crucial component of the “guarding gate“ between the lower and upper reproductive tracts, undergoes significant changes in response to the hormonal cycle. The physical appearance of cervical mucus transitions from dry and pasty (high viscoelasticity) after menstruation to a more slippery, stretchy, and clear (low viscoelasticity) state, facilitating sperm ascent through the cervix to the uterus as estradiol levels rise and ovulation nears [39, 40]. Such changes are associated with an E2-induced increase in the expression of the gel-forming mucin MUC5B [50, 51]. In contrast, the level of MUC5AC in cervical mucus remains constant throughout the menstrual cycle [50, 51]. Furthermore, MUC5B polymerizes into linear bundles with relatively low viscoelasticity, while heavily branched MUC5AC polymerizes into a dense network with a much higher viscoelasticity than MUC5B [41, 45, 47, 80]. Interestingly, we found that GC migrated through (Fig. 6) and aggregated in the high-viscoelastic MUC5AC-dominant mucin hydrogels or mucin protein mixtures more efficiently than the low-viscoelastic MUC5B-dominant mucus or mucins (Figs. 8 and 9). Consistent with our hydrogel results, mucus collected from E2+P4-treated cervical tissue explants promoted GC aggregation more effectively than that collected from E2-treated tissue explants (Fig. 7), even though it remains technically challenging to quantify the molecular ratios of MUC5B versus MUC5AC in the cervical mucus. Furthermore, higher percentages of GC were recovered from inverted, MUC5AC-only hydrogels than from MUC5B-containing hydrogels (Fig. 6E-H). These data together suggest that mucus viscoelasticity is not a limiting factor for GC penetration, but that E2-induced production of MUC5B interferes with GC-GC and/or GC-mucus interactions. However, the mechanisms underlying MUC5B-mediated interference are unknown. GC are known to sialylate their surface utilizing host-produced sialic acid for immune evasion [81, 82], which can further enhance electrostatic repulsion between GC and mucus. Effective migration and strong aggregation of GC in MUC5AC-dominated mucus may explain why GC can remain colonized during menstruation, despite the dramatic shedding of the endometrial lining. The induction of MUC5B by E2 in the proliferative phase potentially improves the mucus’ barrier function against GC.

This study shows that E2 treatment downregulates the expression of CEACAMs at the luminal surface of endocervical epithelial cells (Fig. 3) while enhancing GC colonization, suggesting a CEACAM-independent colonization enhancement. Consistent with this finding, we previously showed that GC colonized epithelial cells at the transformation zone of human cervical explants, which express no or low levels of CEACAMs, and CEACAM-expressing ectocervical epithelial cells equally well [13]. Opa engagement of their human host receptors, CEACAMs, has been shown by multiple groups and various models to enhance GC colonization by inhibiting GC-induced epithelial cell-cell junction disruption and epithelial cell shedding [13, 19]. We have recently shown that GC-induced production of IL-10, a GC colonization enhancer, also requires the downstream signaling of CEACAMs [69]. This CEACAM-independent colonization mechanism remains unknown.

This study examined the impact of female sex hormones on GC infection within a limited time frame (72 h) and concentration window. As an initial study, the treatment concentration and duration were designed to mimic the peak concentration and time only. As a regular menstrual cycle lasts 28 days, graded changes of hormones over 7-10 days may have additional or different effects, which should be carefully examined in future studies. We chose E2 and P4 concentrations, which were much higher than the blood levels, based on previous studies [83–85], considering the local production of these two hormones in the human female reproductive tract, such as the ovary and uterus [86]. However, the hormonal levels at different times of the menstrual cycle at the cervix are not precisely known. Donors of cervical tissue used in this study were younger than 42 years old and at different menstrual cycle phases before surgery, which likely impacts the responses of cervical tissue explants to hormone treatments. To reduce variation, we compared different hormone treatment conditions using cervical tissues from the same donor. Even if individual donors had different baseline hormone levels, the baseline level is the same for tissues from one donor treated with three hormone conditions.

The results presented in this study demonstrate that female sex hormones, E2 and P4, enhance GC colonization at the human endocervix through a CEACAM-independent and mucus-dependent mechanism. Our findings reveal a new mechanism by which GC evade host defense, utilizing the endocervical mucus barrier to enhance colonization.

## Materials and Methods

### Ethics Statement

Human cervical tissue explants were obtained through the National Disease Research Interchange (NDRI, Philadelphia, PA). All tissues were anonymized. The use of human tissues was approved by the Institutional Review Board of the University of Maryland.

### Preparation of hormone-supplemented media

Cervix tissue culture media was prepared using phenol red-free CMRL-1066 media (Islet Media) supplemented with hydrocortisone 21-hemisuccinate sodium (0.1 μg/mL), insulin from bovine pancreas (1 μg/mL), L-glutamine (2 mM), and fetal bovine serum (5%) that had been stripped of hormones overnight with activated charcoal and subsequently filtered. When antibiotics were needed, the media also included penicillin (100 UI/mL) and streptomycin (100 μg/mL). During GC inoculation, antibiotics and hydrocortisone 21-hemisuccinate sodium were omitted from the media [54]. To supplement the media with hormones, estradiol (E2, EMD Millipore) and progesterone (P4, EMD Millipore) were dissolved to a concentration of 1 mM in 100% ethanol, and the ethanol was subsequently evaporated using a SpeedVac. The remaining solid was dissolved in cervix tissue culture media and diluted to the desired concentrations (50 nM E2 and 100 nM P4). An equivalent volume of ethanol was evaporated for use in hormone-free media [61, 84].

### Culture of human cervical tissue explants

Healthy cervical tissues from patients 28-42 years old were received in DMEM with antibiotics through the National Disease Research Interchange (Table 1) within 24 h post-surgery. Tissues were processed and cut into 3∼4 pieces with similar sizes as previously described [13, 54]. Cervical tissue explants were cultured in cervix tissue culture media with or without hormones for 24 h, washed 6 times with antibiotic-free media, and cultured for another 24 h before inoculation with GC.

### Neisseria strains

*N. gonorrhoeae* strain MS11, which expresses phase variable pili and Opa, was grown on GC media plates with 1% Kellogg’s supplement (GCK) for 16-18 h. Using a dissecting microscope, colonies expressing both pili and Opa were visually identified and transferred to a new plate. After 16-18 hours of growth, bacteria were suspended in antibiotic-free cervix tissue culture media containing either no hormone, E2, or E2+P4. The concentration of GC was determined using a spectrophotometer at 650 nm, where an OD of 1 indicated a concentration of 10^9^ GC/mL. GC were diluted in antibiotic-free cervix tissue culture media and inoculated at MOI 10 (10 bacteria to one luminal cervical epithelial cell) for a period of 24 h, with washes 6 and 12 h post-inoculation [13, 54].

### Immunofluorescence analysis of cervical tissue explants

After 24-h inoculation with GC, cervical tissue explants were fixed with 4% paraformaldehyde, perfused with 5% and 15% sucrose, and then 7.5% and 20% porcine gelatin, followed by flash freezing. Frozen tissue samples were cryosectioned, permeabilized, and immunolabeled with antibodies against GC [55] and E-cadherin (BD Biosciences). F-actin and DNA were labeled with phalloidin (Invitrogen) and Hoechst 33342 (Invitrogen), respectively. Tissue sections were imaged using a Zeiss LSM980 confocal microscope, and images were analyzed using NIH ImageJ software.

GC colonization of the endocervix and ectocervix was quantified by the percentage of epithelial cells on the luminal surface that were associated with GC and the fluorescence intensity (FI) of GC staining per µm^2^ of the luminal surface [13]. The data were generated using 10 images per cervix and 3-5 cervixes.

GC penetration in the endocervix was quantified by the percentage of GC-associated cells with GC either at the basal membrane or subepithelium and the percentage of GC staining that was below the basal membrane [13].

GC colony size and density were quantified by outlining GC colonies based on immunostaining and measuring the area of each colony occupied in square microns and the mean fluorescence intensity (MFI) of the gonococcal DNA staining.

Endocervical epithelial cell-cell junction integrity was evaluated by FI ratios of E-cadherin at the cell-cell junction relative to that of the cytosol using FI profiles of lines drawn perpendicular to the cell-cell junction [13]. Epithelial cell shedding was quantified by the percentage of epithelial cells that moved above the endocervical epithelium. Endocervical epithelial cell shape was evaluated by cell height-to-width ratios. The data were generated from 10 randomly acquired images per cervix and ≥2 cervixes.

CEACAM (Santa Cruz) and MUC1 (Santa Cruz) expression in the endocervical epithelium was analyzed by immunostaining and quantified by FI per cell (dividing the total FI from all luminal cells by the number of luminal cells). The data were generated from 5 randomly acquired images per cervix and 3-4 cervixes. Their luminal surface expression was evaluated by the percentage of FI that was localized to the apical surface. MUC1 expression under GC microcolonies was evaluated by ratios of MUC1 FI under a GC microcolony relative to MUC1 FI of an equivalent surface area adjacent to the microcolony. The data were generated from individual GC microcolonies from 5 randomly acquired images per cervix and 3 cervixes.

### Preparation of mucin hydrogels

Mucin hydrogels were generated using a previously published method [80]. Briefly, MUC5B from bovine submaxillary gland (EMD Millipore) and MUC5AC from porcine stomach (EMD Millipore) were rehydrated at a concentration of 4% (w/v) in a buffer (154 mM NaCl, 15 mM NaH_2_PO_4_, 3 mM CaCl_2_, pH 7.4) with stirring for > 2 h. After rehydration, MUC5B and MUC5AC were combined in ratios of 100:0, 75:25, 50:50, 25:75, and 0:100. These mixtures were then allowed to stir for at least 1 h. Separately, 10 kDa polyethylene glycol with a 4-armed thiol functional group (PEG-SH) (Laysan Bio) was dissolved at a concentration of 4% (w/v) in mucin buffer and mixed in a 1:1 ratio with the mucins. The mixture was allowed to stir for 10 min. Following mixing, 20 μL of each mixture was added to the center of a 5 mm diameter rubber O-ring (McMaster-Carr), which had been affixed to a coverslip using vacuum grease. Gels were allowed to solidify for at least 24 h in a humid chamber.

### Bacterial diffusion assay in mucin hydrogels

Piliated MS11 GC were suspended to a concentration of 5×10^3^ GC/mL in antibiotic-free cervix tissue culture media containing 10 μM SYTO9, added to the top of each hydrogel, and incubated for 6 h at 37°C and 5% CO_2_. As imaging controls, goat anti-chicken IgG conjugated to Alexa Fluor 488 (1 µg in 10 µL) were added to the top of hydrogels instead of GC and incubated for 6 h. Following incubation, random Z-stacks were obtained at intervals of 0.6 microns from the bottom of the gel up to 32 µm using a Zeiss LSM980 confocal microscope. Starting from the coverslip, the total FI and MFI of each frame were quantified and plotted against distance from the coverslip.

To determine the number of GC that could be recovered from hydrogels, 100 µL mucin hydrogels in various ratios of MUC5B and MUC5AC were prepared in a 96-well plate and allowed to solidify. Piliated MS11 GC (1.28 x 10^7^ GC in 30 μL antibiotic-free cervical tissue culture media) were loaded on the top of the hydrogel and incubated for 30 min at 37°C and 5% CO_2_. Afterwards, an additional 225 μL of media were added to each well, and the plate was sealed, flipped upside down, and incubated for either 10 min or 2 h. Following incubation, the media was drained, and the hydrogels were collected into 1 mL of media, cut into small pieces, and vortexed to release and disperse bacteria. GC were plated and enumerated.

### Enrichment and quantification of cervical mucus

Cervical tissue explants were hormone-treated and GC-inoculated as described above. The culture media were collected at 6, 12, and 24 h post-inoculation. Post-inoculation supernatants were pooled in equal fractions and treated with protease inhibitor (Sigma-Aldrich). Mucus was then concentrated from supernatants using Amicon Ultra centrifugal filters with a 100 kDa molecular weight cutoff (Millipore) and was stored at –80°C. The mucin concentration in the enriched samples was quantified using a published protocol [87]. Briefly, 50 μL cervical mucin samples in assay buffer (0.01 M Na_2_HPO_4_, 0.04% NaN_3_) were combined with 60 μL of the CNA reagent (200 μL of 0.6 M 2-cyanoacetamide in 1 mL of 0.15 M sodium hydroxide) and boiled for 30 min. The reaction was stopped with the addition of 500 μL of borate (0.6 M, pH 8), and the fluorescence intensity of the solution was measured at an excitation wavelength of 336 nm and an emission wavelength of 383 nm.

### Bacteria aggregation assay

To analyze the effect of sex hormones on GC aggregation, well dispersed piliated MS11 were seeded in 8-well cover glass chambers (2×10^6^ GC in 200 μL) with or without E2 (50 nM) or E2 (50 nM)+P4 (100 nM) and incubated at 37°C and 5% CO_2_ for 6 h before imaging. SYTO9 (10 μM, Invitrogen) was added at 5.5 h post inoculation.

To analyze the effect of cervical mucus on GC aggregation, 0.2% (w/v) cervical mucus in PBS were heated at 65°C for 20 min to kill any microbial contaminants. Well-dispersed piliated MS11GC were stirred with 0.1% mucus for 15 min and seeded in an 8-well cover glass chamber (2×10^6^ GC and 200 μL per well). The chambers were incubated for 6 h at 37°C and 5% CO_2_. SYTO9 (10 μM) was added at 5.5 h.

To analyze the effect of MUC5B and MUC5AC on GC aggregation, MUC5B from bovine submaxillary gland (EMD Millipore) and MUC5AC from porcine stomach (EMD Millipore) were rehydrated at a concentration of 4% (w/v) in PBS with stirring for >2 h. MUC5B and MUC5AC were then combined in ratios of 100:0, 75:25, 50:50, 25:75, and 0:100 and stirred for >1 h. Samples were heated at 65°C for 20 min to kill microbial contaminants. Well-dispersed piliated GC were stirred in a 1:1 mixture with the mucin for 15 min and seeded in an 8-well cover glass chamber (2×10^6^ GC and 2% w/v mucins in 200 μL per well). The chambers were incubated for 6 h at 37°C and 5% CO_2_. SYTO9 (10 μM) was added at 5.5 h.

Random Z-stacks were obtained at intervals of 0.5 microns using a Zeiss LSM980 confocal microscope. Aggregation of GC in the mucus or mucin mixtures was quantified using NIH ImageJ, in which a binary image was created from the sum projections of each Z-stack. This binary image was used to designate regions of interest (ROIs) where GC aggregates were located, and these ROIs were then used for analysis of aggregate number, size, and MFI in the sum intensity projections.

### Western blot analysis of cervical mucins

MUC5B and MUC5AC in mucus enriched from culture media of cervical tissue explants were detected using western blotting as described previously [88]. Mucus (10 µg per lane) were resolved by 4-20% Tris-glycine gel (Invitrogen) under reducing conditions, transferred to a PVDF membrane (Thermo Scientific), and blotted with a monoclonal mouse anti-MUC5B (Santa Cruz) or monoclonal mouse anti-MUC5AC (Abcam) antibody. Mucus (10 µg per lane) in lysates and washes from cultures of human airway epithelial cells (BCi NS-1.1) [89] was used as positive controls.

### Viscoelasticity analysis of animal mucin hydrogels

Viscoelasticity of animal mucin hydrogels was analyzed using multiple particle tracking microrheology as described in a previous study [80]. Briefly, 1 µl of PEG-coated nanoparticles (PEG-NP, 0.002%, w/v) was added to 20 μL of mucins in a vacuum grease-coated O-ring before gelation. Following 24-h gelation, the diffusion of PEG-NP through gels with various MUC5B and MUC5AC concentrations was tracked using a Zeiss Confocal LSM 800 microscope. Multiple 10-sec videos were recorded at 33.3 frames per sec for each sample. Fluorescence microscopy video files were processed using a previously developed MATLAB code capable of tracking multiple particles and calculating the mean squared displacement (MSD) [90, 91].

## Acknowledgments

We thank the UMD CBMG Imaging Core for all microscopy experiments.

